# Potential linkage between *Toxoplasma gondii* Infection and physical education scores of College Students

**DOI:** 10.1101/2020.10.20.346817

**Authors:** Zhijin Sheng, Yu Jin, Yinan Du, Xinlei Yan, Yong Yao

## Abstract

**Objective:** *Toxoplasma gondii* is a worldwide protozoan parasite that could infect virtually all warm-blooded animals, including humans. Our study aimed to investigate the prevalence of *T. gondii* infection in college students at Anhui province, China. Moreover, growing studies demonstrated the association between *T. gondii* infection and host behavioral changes. We also studied the linkage between *T. gondii* and scores of college students.

**Methods:** 2704 serum samples of medical school students attending physical education lessons were collected from September 2017 to September 2019 and evaluated for *T. gondii* IgG antibodies using an enzyme-linked immunosorbent assay (ELISA). We also analysed PE scores of *T. gondii* infected students and *T. gondii* uninfected students.

**Results:** The overall seroprevalence of *T. gondii* was 11.5%. The main risk factors related to *T. gondii* infections were cat in the household and gardening or agriculture activity. Furthermore, in basketball group and football group, scores of *T. gondii* seropositive students were significantly higher than that of seronegative students, while in other sports there is no difference between scores of *T. gondii* infected students and *T. gondii* uninfected students.

**Conclusion:** This is the first report of *T. gondii* seroprevalence in college students in Anhui province, China.

## Introduction

*Toxoplasma gondii* is a kind of intracellular protozoan parasite, which chronically infected about one-third of the world’s population. Humans can be infected by *T. gondii* via three major routes including consumption of unwell-cooked meat or raw meat containing *T. gondii* tissue cysts, ingesting foods or water contaminated with oocysts shed by cats, and congenital transmission to fetus during women pregnancy [1].

In immunocompetent persons, *T. gondii* infection is usually asymptomatic or induces only mild clinical symptoms; however, it may lead to devastating conditions, such as lymphadenitis, meningoencephalitis, or ocular toxoplasmosis in immunosuppressive patients (Transplant recipients, cancer patients, and HIV/AIDS patients). If pregnant women are infected with *T. gondii*, this parasite could cross placental barrier to influence fetal development, resulting in intracranial calcification, mental retardation, chronic chorioretinitis, hydrocephalus, and even fetal death [2].

Though *T. gondii* infection shows no apparent clinical manifestations in immunocompetent people, growing evidence showed that *T. gondii* is associated with increased risk-taking behaviours, in both humans and experimental animals. In humans, latent chronic infection with *T. gondii* has been previously linked with suicidal self-directed violence [3], trait aggression and impulsivity [4], and bipolar disorder [5]. Stefanie K. Johnson *et al*. [6] demonstrated the linkage between *T. gondii* infection and complex human behaviours, including those relevant to business, entrepreneurship and economic productivity. In animals, it has been reported that *Toxoplasma* infection not only reduces a mouse innate aversion to predator odors [7, 8], but also, surprisingly, results in the development of an fatal attraction in rats [9]. The behavioural alterations in *T. gondii* infected rodents may be associated with brain inflammation, because *T. gondii* chronically infects host in the form of tissue cysts in muscle and central nerve system (CNS) [8].

In China’s high education system, physical education (PE) lesson is one optional course, and college students could select sport of their interest, such as football, basketball, and table tennis. Sports need not only physical quality, but also psychological quality. The seroprevalence of *T. gondii* in college students varies from 22.3% in Brazil [10] to 4.8% in USA [11]. Na Yang *et al*. [12] showed that the seroprevalence of *T. gondii* among the newly enrolled undergraduates students in China was 1.63%. However, the impact of *T. gondii* latent infection on study scores of young college students has not been investigated. In present study, we investigated the prevalence of *T. gondii* infection among college students attending PE lessons and analysed the correlation between *T. gondii* infection and scores of participants.

## Materials and Methods

### Participants

A total of 2704 whole blood samples of undergraduates of Anhui Medical University (AHMU) originated from Anhui province of China (between east longitudes of 114°54’ to 119°37’ and north latitudes of 29°41’ to 34°38’) were collected from September 2017 to September 2019 survey the presence of *T. gondii* specific antibodies. The age of the involved students ranged from 20 to 22 years. At AHMU, physical education is optional courses, college students could select which kind of sport they like taking by themselves. In present study, scores of PE of students from six groups (Football, basketball, tennis, table tennis, badminton, and volleyball) were extracted from their annual examination records. Teachers for PE do not know the infection status of students.

### Ethics Statement

This study was approved by the institutional review board at the Anhui Medical University (# 2017USHAEC-026) with written informed consent from all participants. All subjects gave written informed consent in accordance with the Declaration of Helsinki.

### Serological Testing

The sera of all participants were tested for the specific IgG to *T. gondii* using commercial ELISA Kit for IgG Antibody to *Toxoplasma* (Haitai Biomed, Zhuhai, China), according to manufacturer’s instruction. Briefly, the serum sample was diluted with a ratio of 1:100, followed by adding to test well in the antigen-coated plate and incubation at 37 ^°^C for 30 min. After intensive washing for three times with washing solution, 50 μL peroxidase-conjugated anti-human IgG was added to the wells. After 30 min incubation at 37 ^°^C, each well was washed with washing solution for three times. Then, 50 μL “A” solution and 50 μL “B” solution were added to test wells, and the plate was incubated at 37 ^°^C. Ten min later, reaction was stopped by adding 50 μL stopping solution. Microplates were read at an optical density (OD) of 450 nm in the MK3 microplate reader (Thermo Fisher Scientific, Waltham, MA, USA) and ratios (OD 450 value of serum sample/OD 450 value of negative control) were calculated after correction for the OD 450 value of the blank well. The serum samples were considered positive when the ratio was ≥2.1.

### Questionnaire

The questionnaire contained information of basic demographic data, including age, gender, and residence before university. Possible risk factors, including drinking unboiled water, raw or not well-cooked meat (including lamb, beef, pork, fish) and raw vegetable consumption, cat contacts, gardening or agricultural activities and living in urban areas or countryside.

### Statistical Analysis

For the statistical analysis, the SPSS 20.0 software package (IBM, Armonk, NY, United States) was used. Statistical analyses of *T. gondii* prevalence in different variables were performed by χ^2^-test. PE scores of students of seropositive students and seronegative students were analysed using *t*-test. *P*-values less than 0.05 were considered statistically significant.

## Results

### Seroprevalence of *T. gondii* Infection

First, we used ELISA to detect the seroprevalence of *T. gondii* infection among college students attending PE lessons. As shown in Table 1, the overall seroprevalence of *T. gondii* among participants at AHMU was 11.5% (95% CI [10.30-12.70]). There was significant difference in the seroprevalence of *T. gondii* between male students (13.14%, 95% CI [11.51-14.77]) and female students (8.90%, 95% CI [7.17-10.63]) (χ^2^= 10.916, *p*= 0.001). In the univariate analysis, we found two variables associated with anti-*T. gondii* IgG positivity, including cat in the household (*p*= 0.000) and gardening or agricultural activity (*p*= 0.047) (Table 2).

**Table 1.** Seroprevalence of anti-*T. gondii* IgG antibody in 2704 college students at different PE groups in Anhui province, China.

|  | Male |  |  | Female |  |  | Total |  |  |
| --- | --- | --- | --- | --- | --- | --- | --- | --- | --- |
|  | No. positive | No. negative | Positive rate<br>(%, 95% CI) | No. positive | No. negative | Positive rate<br>(%, 95% CI) | No. positive | No. negative | Positive rate<br>(%, 95% CI) |
| Basketball | 75 | 325 | 18.75 (14.91-24.59) | 11 | 106 | 9.40 (4.03-14.77)* | 86 | 431 | 16.63 (13.41-19.86) |
| Football | 64 | 303 | 17.44 (13.54-21.34) | 15 | 123 | 10.87 (5.61-16.13) | 79 | 426 | 15.64 (12.46-18.82) |
| Volleyball | 34 | 212 | 13.82 (9.48-18.16) | 28 | 183 | 13.27 (8.66-17.89) | 62 | 395 | 13.56 (10.42-16.72) |
| Badminton | 17 | 179 | 8.67 (4.70-12.65) | 18 | 192 | 8.57 (4.75-12.39) | 35 | 371 | 8.62 (5.88-11.36) |
| Tennis | 17 | 219 | 7.20 (3.88-10.53) | 9 | 153 | 5.56 (1.99-9.12) | 26 | 372 | 6.53 (4.09-8.97) |
| Table tennis | 11 | 203 | 5.14 (2.16-8.12) | 12 | 195 | 5.80 (2.59-9.00) | 23 | 398 | 5.46 (3.28-7.64) |
| Total | 218 | 1441 | 13.14 (11.51-14.77) | 93 | 952 | 8.90 (7.17-10.63) * | 311 | 2393 | 11.50 (10.30-12.70) |

\*The female vs the male, $p < 0.05$ .

**Table 2.**
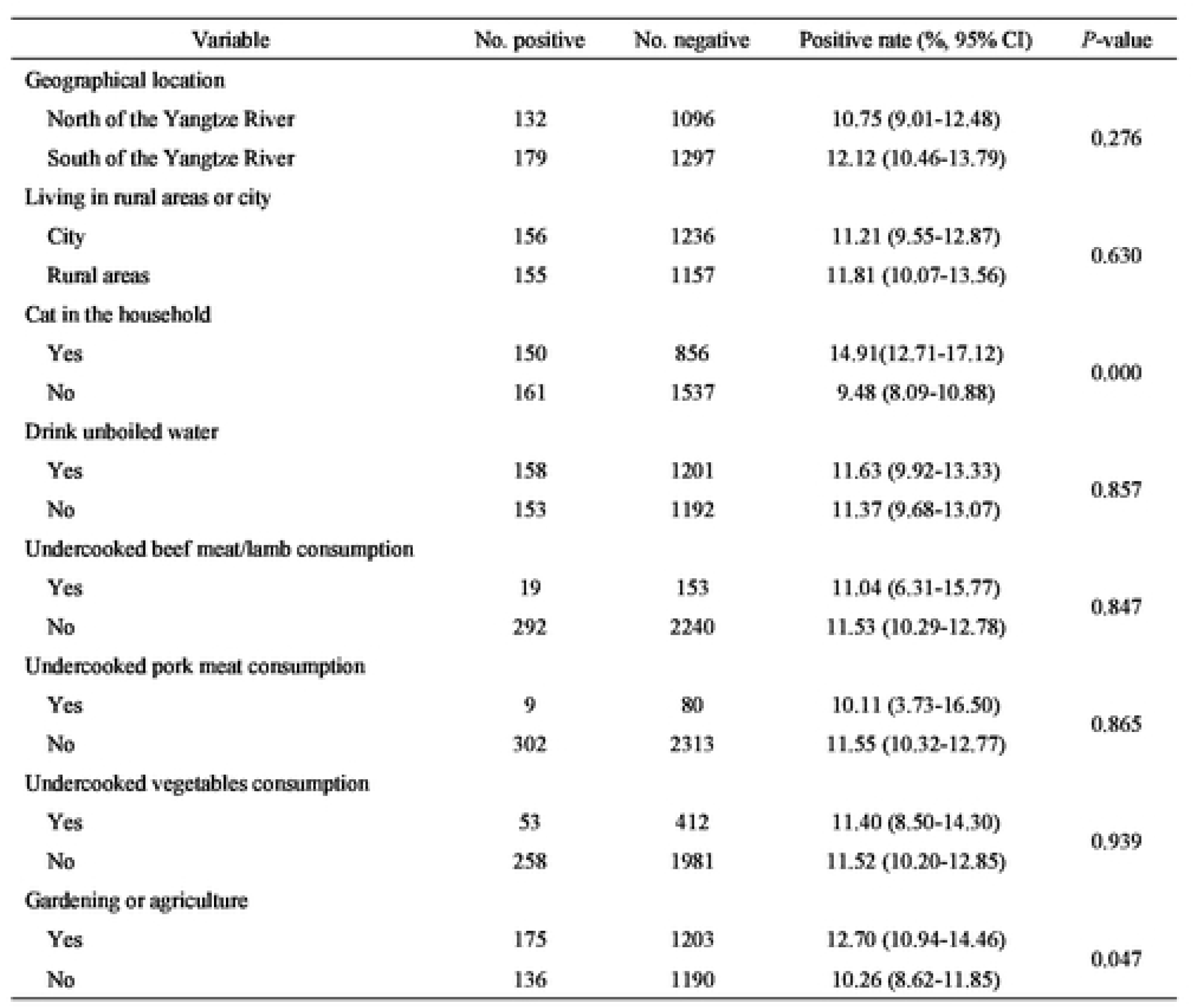
Univariate analysis of the factors associated with *T. gondii* seroprevalence of 2704 college students in Anhui province, China.

| Variable | No. positive | No. negative | Positive rate (%; 95% CI) | *P*-value |
| --- | --- | --- | --- | --- |
| Geographical location |  |  |  |  |
| North of the Yangtze River | 132 | 1096 | 10.75 (9.01-12.48) | 0.276 |
| South of the Yangtze River | 179 | 1297 | 12.12 (10.46-13.79) |  |
| Living in rural areas or city |  |  |  |  |
| City | 156 | 1236 | 11.21 (9.55-12.87) | 0.630 |
| Rural areas | 155 | 1157 | 11.81 (10.07-13.56) |  |
| Cat in the household |  |  |  |  |
| Yes | 150 | 856 | 14.91(12.71-17.12) | 0.000 |
| No | 161 | 1537 | 9.48 (8.09-10.88) |  |
| Drink unboiled water |  |  |  |  |
| Yes | 158 | 1201 | 11.63 (9.92-13.33) | 0.857 |
| No | 153 | 1192 | 11.37 (9.68-13.07) |  |
| Undercooked beef meat/lamb consumption |  |  |  |  |
| Yes | 19 | 153 | 11.04 (6.31-15.77) | 0.847 |
| No | 292 | 2240 | 11.53 (10.29-12.78) |  |
| Undercooked pork meat consumption |  |  |  |  |
| Yes | 9 | 80 | 10.11 (3.73-16.50) | 0.865 |
| No | 302 | 2313 | 11.55 (10.32-12.77) |  |
| Undercooked vegetables consumption |  |  |  |  |
| Yes | 53 | 412 | 11.40 (8.50-14.30) | 0.939 |
| No | 258 | 1981 | 11.52 (10.20-12.85) |  |
| Gardening or agriculture |  |  |  |  |
| Yes | 175 | 1203 | 12.70 (10.94-14.46) | 0.047 |
| No | 136 | 1190 | 10.26 (8.62-11.85) |  |

*T. gondii* seroprevalence among the students for football, tennis, basketball, table tennis, badminton, and volleyball were 15.64%, 6.53%, 16.63%, 5.46 %, 8.62% and 13.56 %, respectively. No difference exists between seroprevalence of male and female students in football, tennis, table tennis, badminton, and volleyball classes, while for students in basketball classes, seropositive rate of *T. gondii* in male students (18.75%, 95% CI [14.91-24.59]) was significantly higher than that of female students (9.40%, 95% CI [4.03-14.77]) (χ^2^= 5.050, *p*= 0.025).

We next analysed whether there was difference in scores of *T. gondii* infected students and *T. gondii* uninfected students. As we can see in Figure 2, in basketball (85.32±3.23 *vs* 83.08±7.19, *p*=0.000) and football (87.00 ± 7.43 *vs* 84.80 ± 6.84, *p*=0.010), scores of *T. gondii* seropositive students were significantly higher than that of seronegative students, while in other sports there is no difference between scores of *T. gondii* infected students and *T. gondii* uninfected students. The similar pattern was observed that scores of seropositive male students were higher than that of seronegative male students, in football group and basketball group (Figure 2 A and B). For female students in basketball group (Figure 2A), seropositive students presented higher scores (85.23±3.13), compared to seronegative students (80.08±8.90).

**Figure 1.**
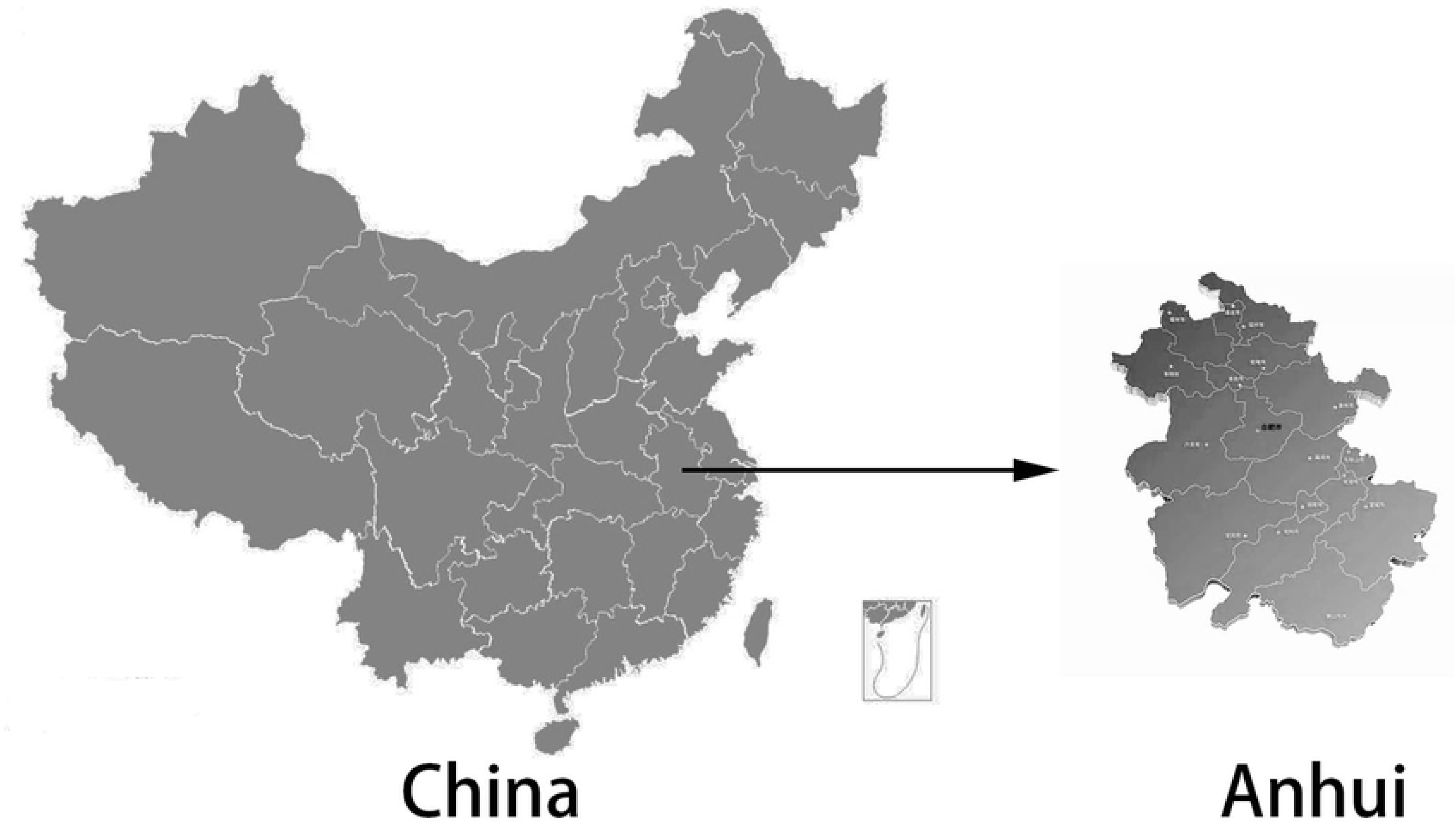
Geographic picture of Anhui province, China.

**Figure 2.**
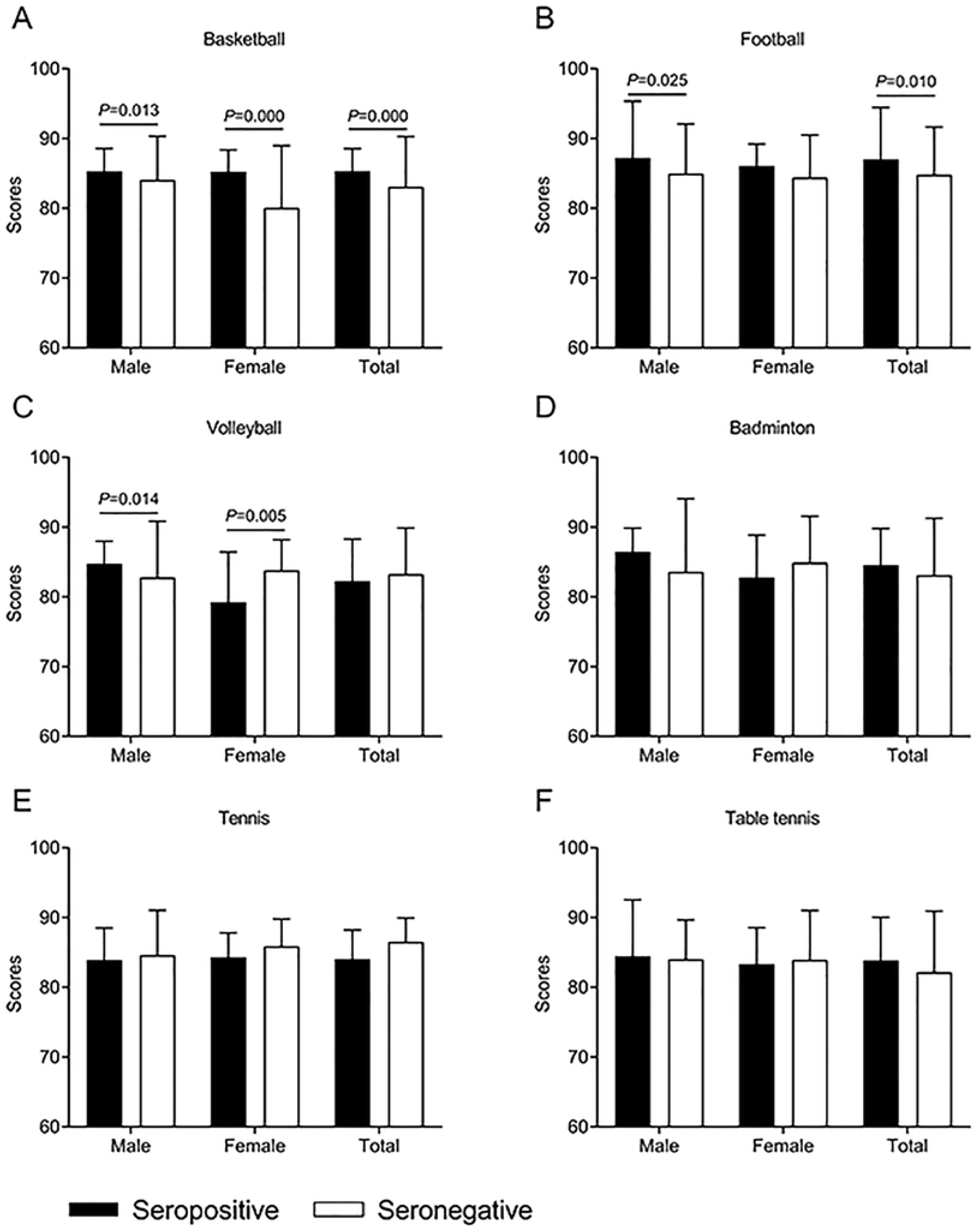
PE scores of 2704 college students in Anhui province, China.

## Discussion

The influence of *T. gondii* chronic infection on host phsyco-behaviours has been a hot point in parasitological study. Our results in present study showed that seroprevalence of *T. gondii* infection of undergraduates attending PE lessons at AHMU was 11.5%. In addition, through analysing PE scores of students from different sport groups, we found that compared with seronegative students, scores of seropositive students in basketball group and football group were apparently higher.

Consistent with previous study showing that the overall anti-*T. gondii* IgG prevalence in China was 12.3% [13], our present study indicated that seropositive rate of *T. gondii* of college student was 11.5%. However, it has been reported that seropositive rate of *T. gondii* in newly enrolled undergraduates from Shenyang Agricultural University (SYAU, China) is 1.63% [12], much lower than our results. This might be due to the different methods used in our study (MAT *vs* ELISA). A study conducted in Henan province (China) using ELISA indicated that the overall seroprevalence of *T. gondii* in primary school children was 9.51% [14]. Furthermore, students in SYAU are enrolled from all provinces around China, whereas students participating in our study are mainly from Anhui province. The primary geographical location of involved student population may contribute to difference in seroprevalence of *T. gondii*. In Korea, 22.5% of children living in the rural areas of Pyin Oo Lwin and Naung Cho, Myanmar were positive for *T. gondii* IgG [15]; In Brazilian, the IFAT method showed a seroprevalence of 22.3% in college students [10]; in Jordan, *T. gondii* IgG antibodies were detected in 66•5% of undergraduate female university students [16]. In addition, like veterinary students having a high prevalence of antibodies to *T. gondii* (5.6%) at Virginia Tech [11], medical students would have a greater chance at exposure to the parasite than an average population of undergraduate students due to increased contact with clinical samples and experimental animals. Though we found that contact with cat (cat in the household) and exposure to soil (gardening or agriculture) were two main factors for college student infection with *T. gondii* in our study, the detailed risk factors for medical students with high seropositive rate of *T. gondii* infection is worth further investigation.

Interestingly, our study found that *T. gondii* chronic infected rates of college students at contact sport groups (basketball and football) were slightly higher than that of non-contact sport groups (badminton, tennis and table tennis). *T. gondii* infection is associated with increased risk-taking behaviours, potentially due to hormonal or neurological changes resulting from parasitic tissue cysts in host CNS [17]. These physiological changes may directly or indirectly enhance infected students to select more challenging sports. We also observed that *T. gondii* seropositive students had higher scores than seronegative students, in basketball group and football group. This result could be partly explained by previous study showing that nations with higher infection rate of *T. gondii* had a lower fraction of respondents citing ‘fear of failure’ in inhibiting new business ventures. In mice and rats, latent *Toxoplasma* infection converted the aversion to feline odors into attraction [18]. A recent report indicates that *T. gondii* lowers general anxiety in infected mice, increases explorative behaviours, and alters predator aversion without selectivity toward felids, which had a positive correlation with the cyst load in host brain [8]. Given that we have no chance to count tissue cysts in human brains, novel methods need to be developed for detection of neuropathological levels of *T. gondii* chronic infected humans.

In conclusion, our study suggests the relatively high seroprevalence of *T. gondii* infection in college students at Anhui province, China. Though we found that PE scores (basketball and football) of seropositive students were higher, the relationship between student PE scores and *T. gondii* infection should be determined by further well-designed investigations.

## Conflict of interest statement

All authors declare that no conflict of interest exists in this study.

## Acknowledgements

Our study is financially supported by Research on the humanities and social sciences of Anhui Province (Grant #: SK2017A0163), Anhui high education quality elevation project (Grant #: 2017jyxm0143), Research Foundation for Universities at Anhui (Grant #: KJ2019A0264) and National Natural Science Foundation of China (Grant #: 31701162), and Program of Inner Mongolia Natural Science Foundation of China (No. 2018BS03015).

